# The structural basis for forkhead box family specificity revealed by the crystal structure of human FOXN1 in complex with DNA

**DOI:** 10.1101/428011

**Authors:** Joseph A. Newman, Hazel Aitkenhead, Angeline Gravard, Ioanna A. Rota, Adam E. Handel, Georg A. Hollander, Opher Gileadi

## Abstract

FOXN1 is a transcription factor that is essential for the development of the thymus and the production of T-lymphocytes. It is a member of a large family of transcription factors which recognize DNA sequences through the conserved Forkhead (FH) domain. FOXN1 recognizes a DNA sequence that is different from the common consensus binding sequence of FH domains, although key binding residues are identical. We present crystal structures of the FH domain of FOXN1, free and DNA-bound, which shed light on the different binding specificities; the structure also revelas the basis of the immunocompromised *nude* mutation, as well as a preferential binding to non-methylated CpG motifs.

**Abstract:** FOXN1 is a member of the forkhead box (FOX) family of transcription factors, and plays an important role in thymic epithelial cell differentiation and function. FOXN1 mutations in humans and mice give rise to the "nude" phenotype which is marked by athymia. FOXN1 belongs to a subset of the FOX family that recognize an alternate consensus sequence (GACGC), which is different from the more widely-recognized canonical sequence consensus RYAAAYA. Here, we present the structure of FOXN1 in complex with DNA at 1.6 Å resolution, in which the DNA sequence is recognised by a mixture of direct and water-mediated contacts provided by residues in an α-helix inserted in the DNA major groove (the recognition helix). Comparisons with other FOX family structures reveal that the canonical and alternate DNA sequences are bound in two distinct modes, with partially different registers for the protein DNA contacts. We identify a single alternate rotamer within the recognition helix itself as an important determinant of DNA specificity, and indicate sequence features in the recognition helix that could be used to predict the specificity of other FOX family members. Finally we demonstrate that FOXN1 has a significantly reduced affinity for DNA containing 5-methylcytosine, which may have implications for the role of FOXN1 in thymic senescence.

## Introduction

The FOX family of transcription factors is one of the largest in humans, with 50 members identified to date(1). FOX family proteins play important roles in various cellular processes including the regulation of cell differentiation, proliferation, metabolism, and senescence. FOX family proteins share a ~100 amino acid DNA-binding domain (Forkhead or FH domain), which is widely conserved throughout evolution from humans to yeast and has been used to classify FOX family proteins into 19 subfamilies (denoted FOXA-FOXS) (2). FOXN1 is a FOX family transcription factor that primarily functions as an activator regulating the development of epithelial cells in skin and thymus. The lack of functional FOXN1 protein expression in humans as well as in other vertebrates causes congenital alopecia universalis, nail dystrophy and athymia, designated the "nude" phenotype (3) (4). The latter feature is due to an arrest in thymic epithelial cell (TEC) differentiation beyond a progenitor cell state and causes severe T cell immunodeficiency. The biological targets of FOXN1 in adult TEC were recently identified in a genome-wide study(5). In addition to the control of genes involved in TEC differentiation and proliferation, this analysis also demonstrated that FOXN1 controls - inter alia - the expression of genes involved in antigen presentation and processing including peptidases, proteasome subunits and protein transporters.

FOXN1 is a 648 amino acid protein with the FH domain located centrally between amino acids 270 and 367. No other recognizable domains have been identified, although the N-terminal region has been implicated in thymic epithelial cell differentiation due to the fact that mice lacking the first 154 amino acids displayed a milder thymus phenotype in comparison the full "nude" phenotype but maintained a normal coat (6). Similarly, an acidic cluster of amino acids within the C-terminal 175 amino acids terminus has been identified to contain the region responsible for transcriptional activation (7).

The canonical recognition motif for forkhead domains has the seven-base FKH consensus pattern RYAAAYA (R= purine and Y= pyrimidine). In contrast, FOXN1 recognizes an alternate 5-bp DNA motif, GACGC. This alternate motif is also bound by a subset of FOX proteins, designated FHL (named after the saccharomyces cerevisiae *Fhl1* gene, the first family member to show this binding property(8, 9). To date, there have been several crystal structures of FH domains bound to the canonical FKH DNA motifs, but none of the alternate FHL motif. The molecular basis of binding to the two very different motifs is not known, and is the subject of this report.

FH domains comprise a subclass of the much larger and more diverse winged helix (WH) superfamily and comprises a single 3 stranded mixed β-sheet flanked on one side by three α helices which form a helix-turn-helix core. The "wings" for which the fold is named are generally less well conserved across family members. The first wing (wing 1) is formed by an extended loop between strands β2 and β3 whilst the second wing (wing 2) constitutes the residues immediately following from strand β3. Early structural studies on human FOXA3/HNF-3 established a conserved mode of DNA recognition whereby the third α-helix (α3 or "recognition helix") is inserted deep within the major groove of the DNA and provides direct and water-mediated sequence-specific contacts to the DNA bases that facilitate DNA recognition(10). To date a number of structures of FOX family proteins have been determined in complex with DNA, including human FOXA2(11), FOXK1(12), FOXO1(13), FOXO3(14), FOXO4(15), FOXP2(16) and FOXM1(17). All structures show the same basic arrangement of direct and water-mediated sequence-specific contacts provided by residues in α3, underlying the recognition of the canonical forkhead (FKH) consensus sequence RYAAAYA.

The recognition of the alternate FHL motif GACGC by FOXN1 and a small subset of forkhead proteins is particularly puzzling because the sequences of the core DNA contacting residues within the recognition helix α3 are strictly conserved even across family members with different specificities. A recent analysis of the evolution of these alternate specificities within the FOX family indicates that the alternate specificity has evolved independently in three different phylogenetic lineages(8). Moreover, some bi-specific proteins have also been identified that are able to bind with high affinity to both motifs. Understanding the basis of recognition of divergent DNA sequences will require molecular structures of FH domain(s) bound to the alternate motif.

In the present study we describe the crystal structures of human FOXN1 both alone and in complex with DNA. This is the first structure of any FOX family member bound to a non-canonical FHL motif GACGC. Detailed analysis of the structure reveals a distinct mechanism used by FOXN1 to recognize its specific DNA motif. Comparisons with previous FOX family DNA complexes show that whilst the conformation of the recognition helix remains largely unchanged, the DNA is bound by FOXN1 in an alternative manner, providing a different register for base specific contacts.

## Materials & Methods

### Cloning Overexpression and Purification

FOXN1 constructs corresponding to the FH domain (270–366) and full length (1–648) were cloned in the vector pNIC28-Bsa4 using ligation independent cloning and transformed into *E. coli* BL21(DE3)-R3-pRARE2 cells for overexpression(18). Cells were grown at 37 °C in TB medium supplemented with 50 ug/ml Kanymycin until an optical density of 2–3 and induced by the addition of 0.3 mM IPTG and incubated overnight at 18 °C. Cells were harvested by centrifugation. For purification, cell pellets were thawed and resuspended in buffer A (50 mM HEPES pH 7.5, 500 mM NaCl, 5% glycerol, 10 mM imidazole, 0.5 mM Tris (2-carboxyethyl) phosphine (TCEP)), with the addition of 1x protease inhibitor set VII (Merck, Darmstadt, Germany). Cells were lysed by sonication and cell debris pelleted by centrifugation. Lysates were loaded on to a Ni-sepharose IMAC gravity flow column (GE healthcare), washed with 2 column volumes of wash buffer (buffer A supplemented with 45 mM imidazole), and eluted with 300 mM imidazole in buffer A. The purification tag was cleaved with the addition of 1:20 mass ratio of His-tagged TEV protease during overnight dialysis into buffer A. TEV was removed by IMAC column rebinding and final protein purification was performed by size exclusion chromatography using a HiLoad 16/60 Superdex s75 column in buffer A. Protein concentrations were determined by measurement at 280nm (Nanodrop) using the calculated molecular mass and extinction coefficients. Protein masses were checked by LC/ESI-TOF mass spectrometry, which discovered the intact mass of the full length construct of 65540 Da corresponding to a single truncation at residue 614 in the C-terminus. This construct is therefore referred to as 1–614 in the main text.

### Crystallization and Structure Determination

For crystallization the forkhead domain construct was concentrated to 10 mg/ml using a 10,000 mwco centrifugal concentrator and buffer exchanged to 10 mM Hepes pH 7.5, 250 mM NaCl, 0.5 mM TCEP. DNA was prepared by mixing the oligos 5'-GGTGGCGTCTTCA and 5'-TGAAGACGCCACC in a 1:1 ratio at a concentration of 500 μM, heating for 5 minutes at 94 °C, and letting cool slowly on a heat block. DNA and protein were mixed in a 1.2:1 molar ratio with final protein concentration of 5mg/ml. The protein:DNA complex crystals grew from conditions containing 8% PEG4000, 0.1M acetate pH 4.5, and the DNA-free crystals grew from conditions containing 10% ethylene glycol, 0.25M potassium citrate tribasic, 32% PEG3350. Both crystals were cryo-protected by transferring to a solution of mother liquor supplemented with 25 % ethylene glycol and flash-cooled in liquid nitrogen.

Data were collected at Diamond Light Source beamline I04 (DNA complex) and I03 (DNA-free). Diffraction data were processed with the programs DIALS(19) (DNA complex) and XDS(20) (DNA free), and the structures were solved by molecular replacement using the program PHASER(21) with the FOXK1-DNA complex(12) structure as a starting model. Model building and real space refinement were performed in COOT(22) and the structures refined using PHENIX REFINE(23). The structural coordinates and structure factors were deposited in the Protein Data Bank, PDB: 5OCN (DNA free) and 6EL8 (DNA complex). A summary of the data collection and refinement statistics is shown in Table I.

**Table 1.**
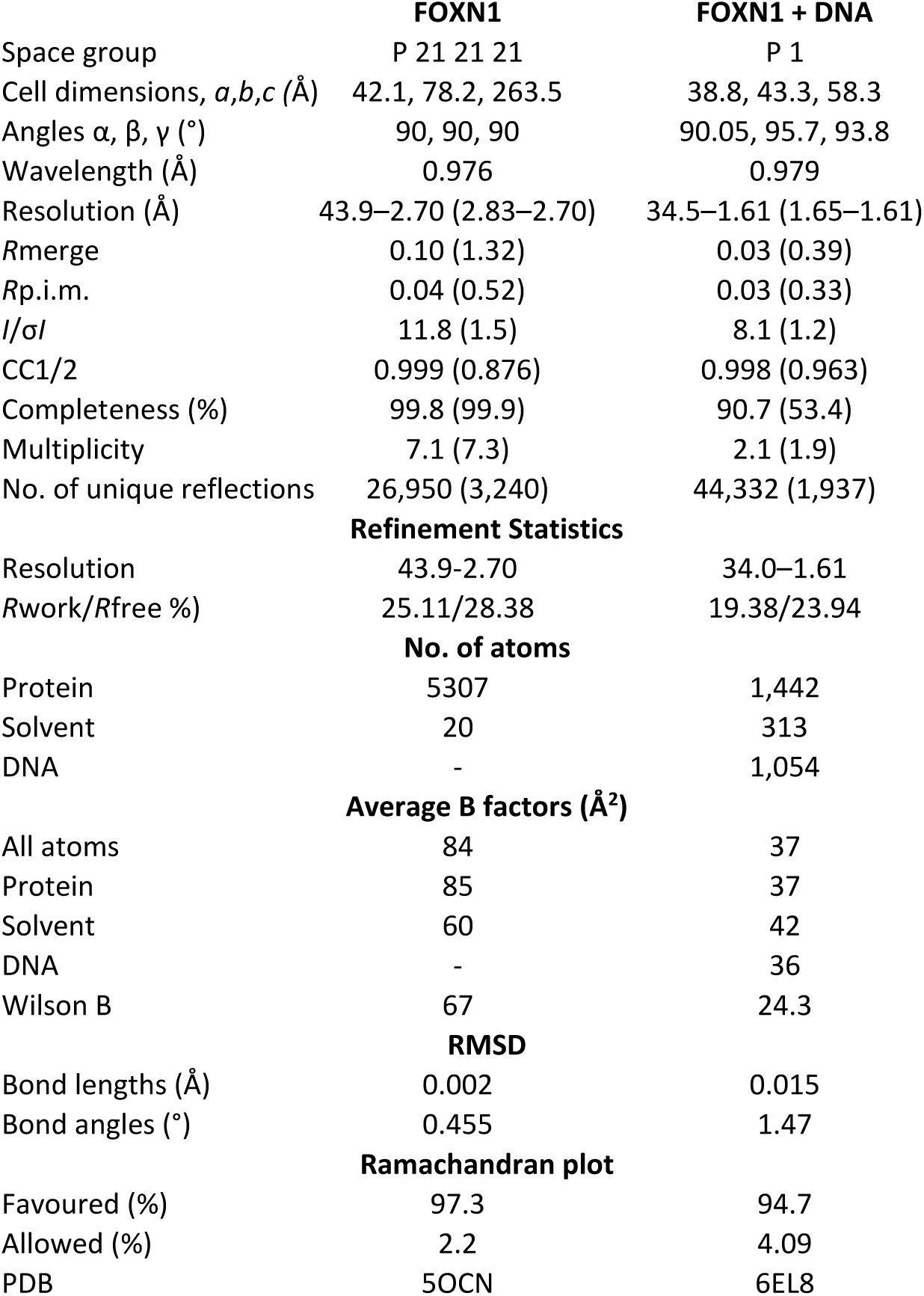
Data collection and refinement statistics

|  | **FOXN1** | **FOXN1 + DNA** |
| --- | --- | --- |
| Space group | P 21 21 21 | P 1 |
| Cell dimensions, *a,b,c* (Å) | 42.1, 78.2, 263.5 | 38.8, 43.3, 58.3 |
| Angles $\alpha, \beta, \gamma$ (°) | 90, 90, 90 | 90.05, 95.7, 93.8 |
| Wavelength (Å) | 0.976 | 0.979 |
| Resolution (Å) | 43.9–2.70 (2.83–2.70) | 34.5–1.61 (1.65–1.61) |
| *R*merge | 0.10 (1.32) | 0.03 (0.39) |
| *R*<sub>p.i.m.</sub> | 0.04 (0.52) | 0.03 (0.33) |
| *I*/σ*I* | 11.8 (1.5) | 8.1 (1.2) |
| CC1/2 | 0.999 (0.876) | 0.998 (0.963) |
| Completeness (%) | 99.8 (99.9) | 90.7 (53.4) |
| Multiplicity | 7.1 (7.3) | 2.1 (1.9) |
| No. of unique reflections | 26,950 (3,240) | 44,332 (1,937) |
| **Refinement Statistics** |  |  |
| Resolution | 43.9–2.70 | 34.0–1.61 |
| *R*work/*R*free (%) | 25.11/28.38 | 19.38/23.94 |
| **No. of atoms** |  |  |
| Protein | 5307 | 1,442 |
| Solvent | 20 | 313 |
| DNA | - | 1,054 |
| **Average B factors (Å<sup>2</sup>)** |  |  |
| All atoms | 84 | 37 |
| Protein | 85 | 37 |
| Solvent | 60 | 42 |
| DNA | - | 36 |
| Wilson B | 67 | 24.3 |
| **RMSD** |  |  |
| Bond lengths (Å) | 0.002 | 0.015 |
| Bond angles (°) | 0.455 | 1.47 |
| **Ramachandran plot** |  |  |
| Favoured (%) | 97.3 | 94.7 |
| Allowed (%) | 2.2 | 4.09 |
| PDB | 5OCN | 6EL8 |

### Electrophoretic mobility shift DNA binding assays

DNA binding was measured using an Electrophoretic mobility shift assay. The probes consisted of the following oligonucleotide sequences annealed to their complementary strands (the FOXN1 consensus sites are in bold): 5'- GCAGCA**GACGC**AACAGAGCGA**GACGC**CAGGG, and 5'-GCAGCA**GAC^ME^GC**AACAGAGCGA**GAC^ME^GC**CAGGG for the methylated DNA probe. Radiolabeled double-stranded DNA probes were prepared by incubating the forward strand oligonucleotides for 2 h at 37 °C with T4 polynucleotide kinase in the presence of [γ-^32^P]ATP. Complementary (non-radiolabeled) oligonucleotides were added in a 2-fold excess, and the mixture was heated to 95 °C and allowed to cool slowly to room temperature. The double-stranded DNA probes were purified on a Bio-Rad P6 micro-biospin column equilibrated in 10 mM Tris pH 7.5, 50 mM NaCl. EMSAs were performed by incubating radiolabeled probe (at a concentrations raging between 0.1 nM and 5 nM depending on the age of the probe) with a 2-fold serial dilution of FOXN1 construct 1–614. The final reaction volume was 7ul (5ul protein stock, + 2ul probe) with a final buffer containing 50 mM Tris-HCl, pH 7.5, 25 mM NaCl, 50 mM L-arginine-HCl, pH 7.5, 0.5 mM EDTA, 0.1% Tween 20, 2 mM DTT, and 5% glycerol. Reactions were incubated for 30 min at room temperature, and 4 μl of each reaction was loaded on to a 12% native polyacrylamide gel electrophoresis in Tris borate EDTA buffer. Gels were ran at 170 V for 70 minutes on ice. Gels were visualized using phosphorimaging, and quantitation was performed using quantity one 1-D analysis software (Bio-Rad). Apparent dissociation constants were calculated using a sigmoidal four-parameter logistic nonlinear regression model in PRISM (GraphPad).

### FOXN1 ChIP-seq enrichment analysis

FOXN1 ChIP-seq data was derived from ref (5). The location of CpG islands was obtained from the UCSC Genome Browser (mm10 genome build). Enrichment was assessed using GAT with 10,000 permutations and a background determined by H3K27ac peaks in cortical thymic epithelial cells(24).

## Results & Discussion

### Structure of human FOXN1

The structure of the forkhead domain (residues 270–366) of human FOXN1 was determined in the presence and absence of DNA at 1.6 Å and 2.7 Å resolution, respectively. The electron density was of overall good quality with a single seven residue loop and the final 5 residues at the C-terminus not visible in the electron density maps, presumably due to disorder. A summary of the data collection and refinement statistics are shown in Table I. Significant conformational changes of FOXN1 were not apparent upon the molecule's binding to DNA (rmsd of 0.7 Å over 84 residues). The overall structure of FOXN1 is also very similar to a number of other FH domain family proteins such as FOXM1(17), FOXP2(16) and FOXK1(12) (~ 1.1 Å rmsd over ~80 residues), despite an only modest pairwise sequence identity of approximately 40 %. The most prominent differences between the various structures lie in the sequence, length and conformation of the wings. Wing 1 of FOXN1 constitutes a relatively long loop that is partially disordered at one end, regardless of whether the molecule is unbound or complexed to DNA (Figure 1A). The Wing 2 region forms an additional a-helix (α4) which packs against the rest of the helical core and ends with a stretch of positively charged residues that also remain disordered in electron density maps. Another feature that varies among forkhead structures is the conformation of the loop between helices α2 and α3, which in FOXN1 forms an additional short 3-_10_ helix.

**Figure 1.**
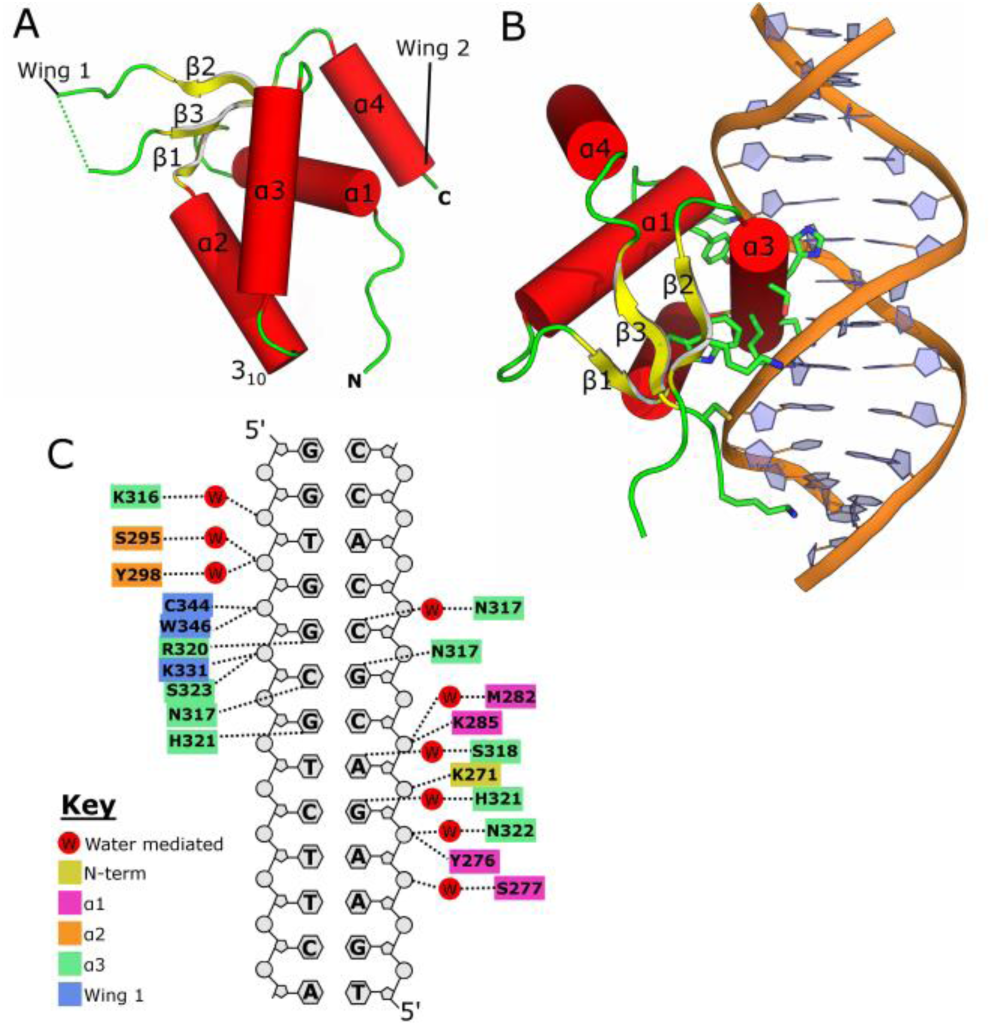
**-** Structure of FOXN1 and the FOXN1 DNA complex. (**A**) Overall structure of FOXN1 with secondary structure elements labelled. (**B**) Structure of the FOXN1 DNA complex with key interacting residues shown in the stick format. (**C**) Schematic view of the FOXN1 DNA interaction with polar contacts marked.

### Recognition of specific DNA sequences by FOXN1

The consensus sequence for FOXN1 binding has been identified as an invariant stretch of 5 residues with a consensus sequence 5'-GACGC(5, 25). For the determination of the DNA complex structure, we used a specific 13-nucleotide double-stranded DNA sequence that contained a single copy of this motif flanked by sequences derived from the mouse *Psmb11* promoter (a high confidence target of mouse FOXN1 encoding the proteasome component β5t; TGAA**GACGC**CACC). Comparable to other winged-helix superfamily proteins, the third helix of FOXN1 (also referred to as the recognition helix) inserts deep into the major groove of the DNA and provides specific contacts to the nucleotide bases (Figure 1B). Other regions contributing towards DNA binding include the N-terminus, the start of the al helix and several residues within and flanking wing 1 that contact the phosphodiester backbone via direct or water-mediated polar interactions (Figure 1C). The overall conformation of the DNA is a modified B form with slight bending towards the protein and concomitant widening of the major groove to accommodate the insertion of α3.

The pattern of polar interactions in the DNA complex structure are summarized schematically in Fig 1C; the details of key protein:base interactions are shown in Fig 2. FOXN1 recognizes the first base pair (G-C) of the GACGC motif primarily by water-mediated contacts, with two waters within hydrogen bonding distance of H321 making polar contacts to the N7 and O6 groups on the Guanine base (Figure 2A). It is not clear how this interaction can underlie a unique discrimination of a G-C pair as the two water molecules could presumably also make hydrogen bonds to an Adenine, whilst the nearby histidine (H321) could be either a hydrogen bond donor or acceptor depending on the protonation state of the NE2 nitrogen. Thus, some degree of indirect motif recognition may play a role at this position. The second base pair (A-T) is also close to a network of highly coordinated waters, with a single water (Water 128) making a pair of hydrogen bonds to the Adenine N7 and N6. Although the pattern of hydrogen bond donors and acceptors within the water network would permit both donor and acceptor at the N6 positon (consistent with the binding of A or G at this position), the angles appear to be more favourable for the water to accept a hydrogen bond from the N6 position (135° versus 64°), thus favouring Adenine at this position (Figure 2B). Further direct contacts are provided by the side chains of S318, which is in a position to donate a hydrogen bond to the Adenine N7, and H321 which makes van der Waals contacts to both the O4 and the (methyl) C7 on the corresponding Thymine. The third base pair (C-G) is recognized by FOXN1 via a direct hydrogen bond to the Guanine N7 donated by the ND1 of H321, together with close van der Waals contacts to the Cytosine C5 which, if replaced by a Thymine, would cause steric clashes with the side chain of S318 and the main chain carbonyl of G314 (Figure 2C). Given that this position forms one of two CpG sites on the motif, we also expect that FOXN1 would, by the same mechanism, be unable to bind to 5-methylcytosine at this position. The fourth base pair (G-C) lies very close to and is directly under the path of the recognition helix, forming two hydrogen bonds with N317: the N4 of the Cytosine donates a hydrogen bond to the main chain carbonyl, and the O6 of the Guanine accepts a hydrogen bond from the side chain ND2 (Figure 2D). Similarly to the case at position 3, the C5 of the Cytosine at position 4 makes close contacts with the C-terminal half of the recognition helix and severe steric clashes would occur by the presence of a 5-methyl group (Thymine or 5-methycytosine). Finally the fifth base pair (C-G) is recognized directly by FOXN1 through a pair of hydrogen bonds donated by R320 to the N7 and O6 of the guanine (Figure 2E). This mode of recognition of a guanine by an arginine side chain is common to several other families of transcription factors. The central importance of this interaction is reflected by the fact that a single point mutation at position R320 in either human or mice results in a loss of FOXN1 transcriptional activity and consequently in the "nude" phenotype (26, 27).

**Figure 2.**
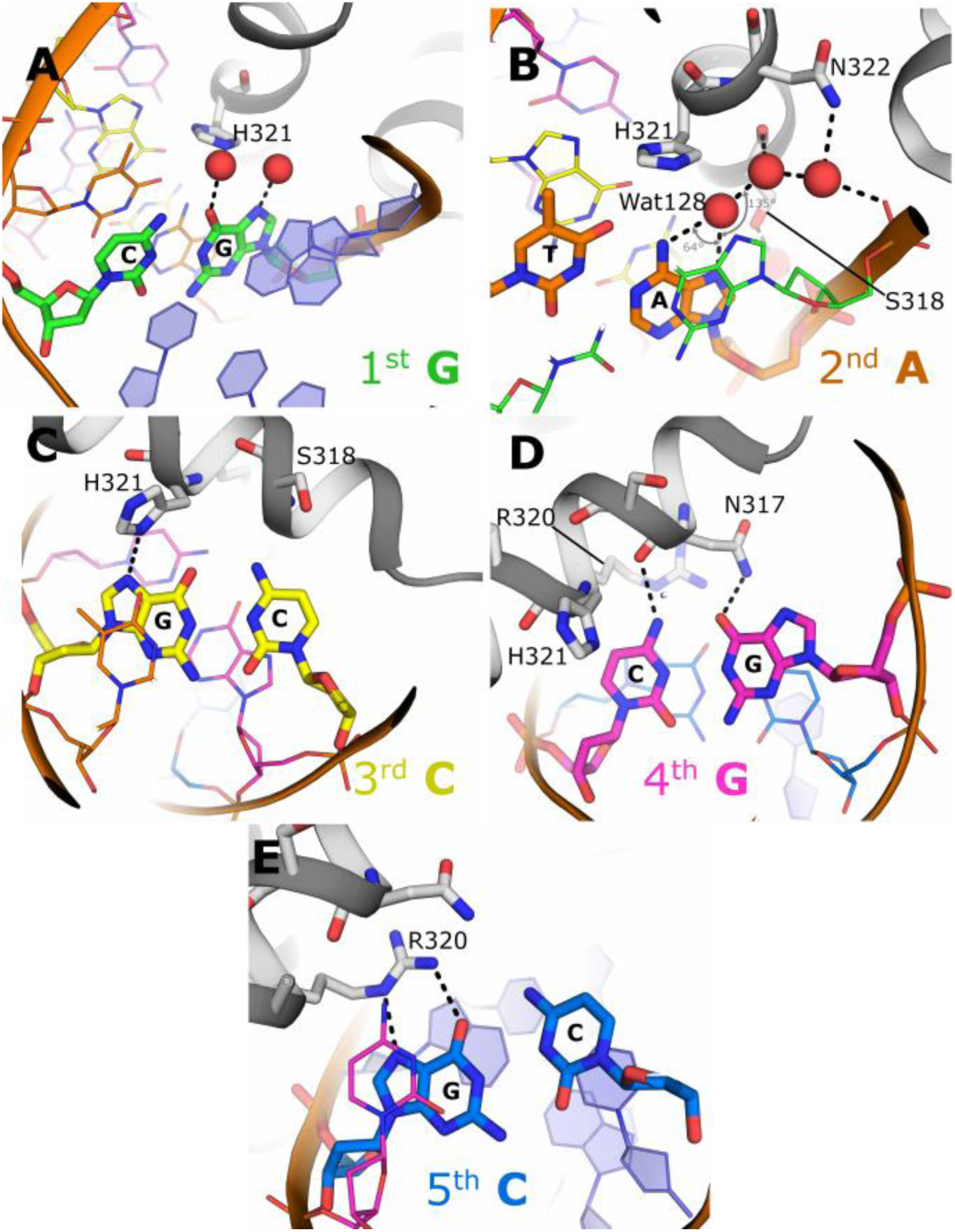
Details of the recognition of the alternate forkhead motif GACGC by FOXN1. Key interactions that determine substrate specificity at the 1^st^ to 5^th^ positions of the motif are shown in panels **A-E** respectively, interacting residues are shown as stick format and water molecules are shown as red spheres.

### Comparison of FKH and FHL recognition modes

We can now address the question of how FH proteins bearing the same recognition helix sequence are able to bind and recognise two distinctly different DNA motifs; the 7-nucleotide canonical FKH consensus sequence RYAAAYA, and the alternate 5-nucleotide FHL sequence GACGC. Comparing the FOXN1 mode of recognition detailed above with previous structural studies on FOX family proteins demonstrates both similarities and differences in the DNA:protein interface, which provide insights into how this duality may be achieved. Contrary to expectation, both the general positioning within the DNA major groove, and the rotamers of key DNA contacting side chains within the recognition helix are generally well conserved with one notable exception: FOXN1's side chain of N317 points towards the N-terminus of α3 whereas the corresponding side chain of other FOX family structures point in the opposite direction, making extensive contacts with the DNA bases (Figure 3A). Notably, in the DNA-free FOXN1 structure this same residue is in the same conformation as in the DNA-bound structures of other FOX proteins, indicating an induced fit in the FOXN1-DNA complex. The structures of several FKH DNA complexes have been determined in various configurations including that of FOXO3A with a single FKH motif(14), FOXK1 in complex with a tandem repeat(12), and FOXM1 in complex with an inverted repeat(17). Although there are specific and important differences in the exact sequence that each one of these FOX family members recognizes, and in the various regulatory mechanisms that determine binding affinity, collectively these molecules share an overall mechanism of base recognition. The various mechanisms are described in more detail in the respective references but will be summarised here to highlight the similarities and differences with DNA recognition by FOXN1.

**Figure 3.**
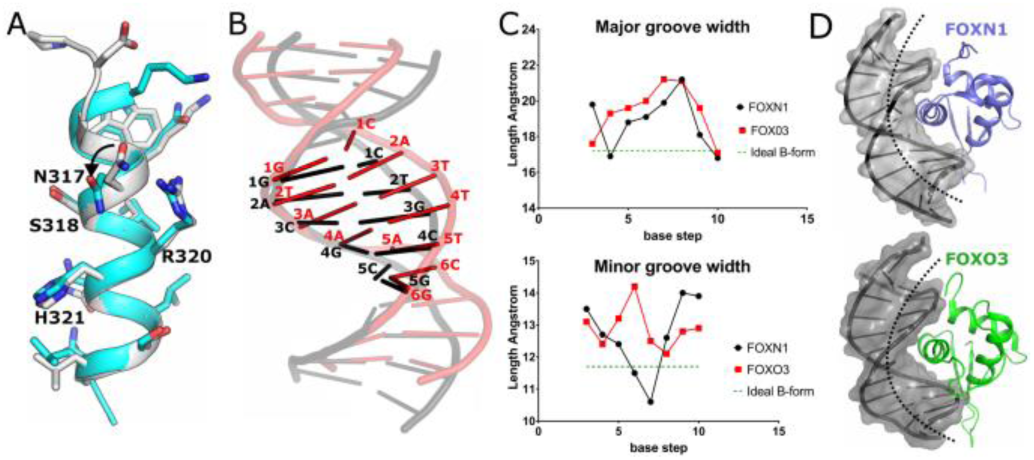
Comparison of the mode of recognition of the FHK and FHL consensus sequences. (**A**) Comparison of the recognition helix of the FHL specific FOXN1 (shown in grey) with the FHK specific FOXK1 (shown in cyan). (**B**) Comparison of the DNA conformations of the FHL site bound to FOXN1 (black) and the FKH site bound to FOXO3 (red), DNA molecules are superposed based on the structural alignment of the FOXN1 and FOXO3 proteins, with the respective motifs highlighted. (**C**) Comparison of the major and minor groove widths for FOXN1 and FOXO3 with values for ideal B-form DNA shown for reference. (**D**) Space filling representation of the DNA bound to FOXN1 (upper panel) and FOXO3 (lower panel) the more prominent bending and greater distance to the recognition helix can be seen in the FOXO3 DNA complex structure.

The most dramatic difference between the DNA complexed to FOXN1 and that to other FOX proteins is a change in the register of the DNA: the recognition helix of FOXN1 interacts with a 5-nucleotide FKH sequence GACGC, while that helix in other FOX proteins interacts with a 6-bp stretch (RYAAAY). This is achieved by the intercalation of the second (T) nucleotide between the positions occupied by nucleotides 1 and 2 in the FKH motif, without extending the physical length of the DNA motif. Some of the same residues in the recognition helix are used in both binding modes, in particular the binding of H321 and R320 (or their equivalent). This change in register is accompanied by dramatic changes in the conformation of the DNA. Most prominent is the change in the inclination of the bases with respect to the helix axis (up to 24° in the FOXO3 DNA structure compared to near-zero in FOXN1 (Figure 3B and table S1). This alteration is sufficient to squeeze the 6 bases of the RYAAAY FKH motif into the same helical rise as the 5-base GACGC FHL sequence. Other less striking differences included high positive roll angles which mainly occur in TA or TG dinucleotide steps in the FKH DNA and significant base pair opening, with generally negative slide and positive roll angles (a full list of DNA geometrical parameters is shown in Table S1). In general, the DNA in the FOXN1 structure was closer to canonical B-form DNA than the FKH DNA structures of other FOX members, although in both cases there is some widening of the major groove (Figure 3C). Taken together, these effects give rise to a more prominent bending of the DNA towards the protein in the FKH complexes, placing the recognition helix generally more distant from the nucleobases (Figure 3D) (i.e. around 2Å further away at the 5' end of the motif). This increased distance means that residues of the recognition helix that make direct contacts with the bases in the FOXN1 structure engage in different, water-mediated contacts with bases in the FKH complexes.

A detailed comparative analysis of the base contacts for each of the two binding modes is shown in Figure 4. The first base of the FKH motif is either an Adenine or Guanine, which is contacted by two water molecules coordinated by an adjacent histidine, similar to the interaction of FOXN1 with the first Guanine of the FHL sequence; as discussed above, this interaction cannot distinguish between A and G. The second base pair (A-T) in the FKH motif is located between the first and second positions in the FOXN1 structure and does not have an equivalent in that structure. The mode of recognition of the second base of the FKH motif is difficult to determine, although there are possible van der Waals contacts with the Thymine methyl. At the third position an invariant Adenine base is recognized by a water network that is similar to that used by FOXN1 to recognize Adenine at position two of the FHL sequence (Figure 4, bottom left panel). The fourth base pair of the FKH motif is also an invariant A-T and is recognized by a combination of a hydrogen bond donated by the equivalent to H321 to the complementary Thymine O4 and a bidentate hydrogen bond via the equivalent of N317, which can recognize uniquely the donor-acceptor pair on the Adenine. In the FOXN1 structure the latter interaction does not occur due to the alternate rotamer of N317, and H321 is hydrogen bonded to the Guanine N7 which occupies approximately the same position as the Thymine O4 (Figure 4, top right). The fifth base pair in FKL (an invariant A-T) is recognised by a combination of water mediated contacts to the complementary Thymine O4 and hydrophobic interactions with the Thymine methyl. In contrast, the equivalent in positon in FOXN1 is a Guanine-Cytosine pair which is recognized by direct rather than water-mediated hydrogen bonds from N317 and close contacts to the cytosine C5 position (Figure 4, centre right). Finally the sixth position of the FKH motif is commonly Cytosine although Thymine is also possible. In both FOXN1 (positon 5) and FKH recognition this is achieved via hydrogen bonds from R320 to the complementary Guanine or Adenine, although in the latter the distance between the guanidinium group and nucleobase is too far to make a bidentate interaction, hence the more relaxed specificity (Figure 4, bottom right).

**Figure 4.**
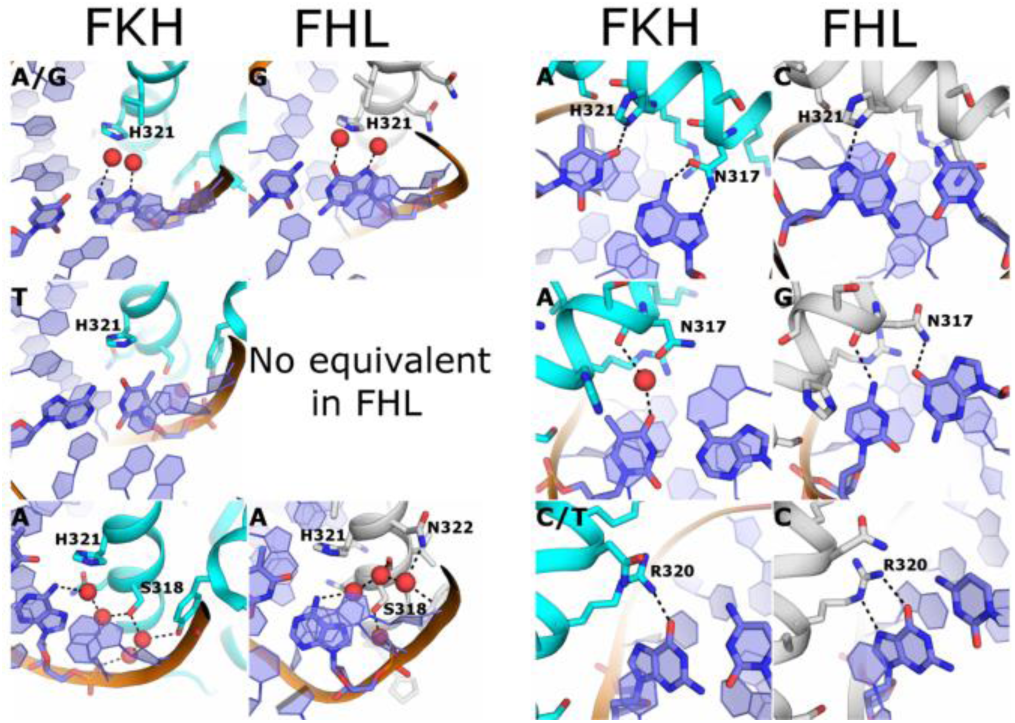
Comparison of the details of the recognition modes of FKH (left hand column) and FHL (right hand column) motif binding. At each position in the respective motifs the contacts to the protein that mediate recognition are show, with the consensus sequence shown throughout in the top left hand corner.

In summary, the interactions which recognize the first, third and sixth positions of the FKH motif are analogous to the first, second and fifth positions respectively the FHL motif, whilst the alternate roamer of N317 switches from recognizing the Adenine at position 4 in the FKH motif to the Guanine at position 4 of the FHL motif. Hydrophobic contacts to Thymine methyl groups have been found to be important for FKH DNA binding in FOXO3(14), whereas in the FOXN1 close contacts to the Cytosine C5 appear to actively preclude binding of Thymine at these positions.

### Structural determinants of FHL binding

A comprehensive analysis of FOX family protein binding specificities established three distinct subgroups of DNA specificities(8), firstly the FKH-specific proteins which comprise by far the largest and most varied group (indicating a FKH specific ancestral origin), a small number of bispecific proteins such as human FOXM1 and FOXN2/3 and finally the FHL-specific groups which include human FOXN1/4 and fungal Fox3. As detailed above, one requirement for FHL binding is the alternative rotamer of N317. Looking at the surrounding context of this residue in both classes of structures, a clear difference can be seen in the conformation of the N-terminus of the recognition helix. Specifically, some of the FKH-specific proteins have an additional turn of the α-helix with a hydrogen bond formed between the main chain amide of the equivalent to N317 and the carbonyl of the -4 residue. This contact appears to prevent the alternate rotamer of the asparagine due to a steric clash (Figures 3A, 5A). Looking at the sequences in this region, a pattern can be recognized in which the FHL-specific proteins have a proline followed by a negatively charged aspartate and a glycine with the general motif PDGW (Figure 5B). The glycine residue is situated at the helix to coil transition immediately above N317, and occupies a region of the Ramachandran plot specific to glycine. The negatively charged aspartate is shifted significantly (3.5 Å) away from the equivalent residue in the FKH binding structures and would have appeared to make unfavourable charge interactions with the DNA backbone if placed in this position. In contrast, the FKH binding family members generally have either a glutamine or a positively charged residue at this position, which in the structures of FOXK1(12) and FOXA2(11) make polar contacts to nearby DNA backbone phosphates, presumably further stabilizing the extended α-helical conformation. Whilst the glycine at position 314 in FOXN1 appears to be a requirement for FHL binding, it may not be sufficient to confer this activity, since a number of apparent FKH specific proteins also contain a glycine at this position (Figure 5B), although in many cases the specificities of these proteins are inferred from sequence homologies and are not actually measured, and may thus be different to what is presumed. The features that distinguish FHL-binding FOX family members from the bispecific variants are less clear, since the DNA-free FOXN1 structure also contained the same rotamer as found in the FKH binding structures. We propose that the fact that bispecific proteins such as FOXM1 contain the flexible glycine residue that allows the FOXN1-like rotamer of N317, while also maintaining a DNA phosphate contact which is absent in FOXN1 (17), accounts for the dual binding specificity.

**Figure 5.**
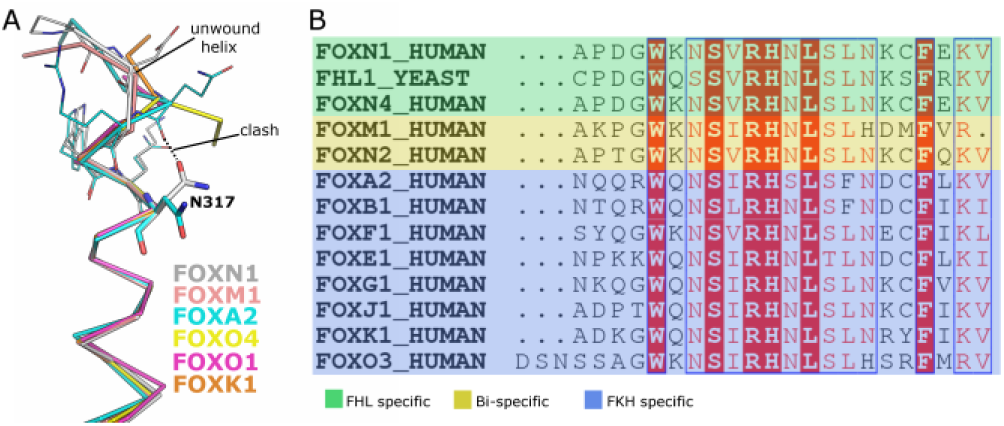
Sequence determinants of FHL versus FKH specific binding. (**A**) Structural alignment of the N-terminal regions of the recognition helix with the alternate rotamers of N317 shown. The extended helix present in a subset of the FKH DNA complex structures would prevent the alternate roamer due to steric clashes. Multiple sequence alignment of a subset of the FOX family proteins from humans and yeast. The specificities of the various proteins were either measured directly, or inferred from the phylogenetic analysis in (17).

### Affinity of FOXN1 for normal and methylated DNA

One further prediction to emerge from the FOXN1 DNA complex structure is that methylation of the CpG within the FHL motif may significantly reduce FOXN1 binding. We have tested this hypothesis using a FOXN1 construct spanning residues 1–614 (we were unable to purify full length FOXN1 due to proteolysis of the C-terminus), and a high-confidence FOXN1 binding site found within the promoter of the proteasome subunit PSMA7 containing two copies of the FHL motif in a tandem arrangement. As can be seen in Figure 6, the non-methylated DNA was bound relatively tightly; while there may be to be multiple shifted species of different electrophoretic mobilities, an apparent Kd of around 100 nM could be calculated from the data. In contrast, DNA containing 5-methylcytosine at both strands of the CpG sites within the FHL motifs was bound significantly less tightly, with an apparent Kd of 1.3 μM. This >10 fold difference in affinity may target FOXN1 binding to CpG islands, regions of non-methylated DNA that are known to occupy the promoter regions of most human genes(26). Indeed analysis of the PSMA7 promoter reveals a very strong prediction for a CpG island (>1000 nucleotides with a GC content of >70%, and a ratio of observed to expected CpG dinucleotides of around 1.0).

**Figure 6.**
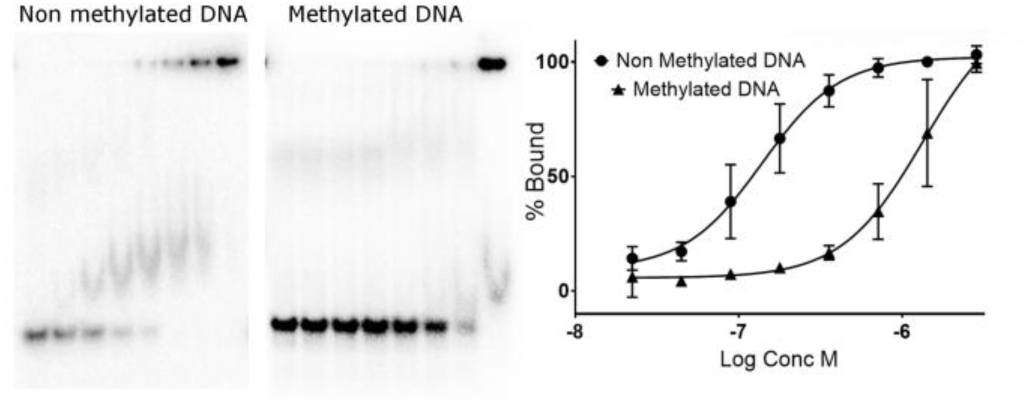
Electrophoretic mobility shift assay showing FOXN1 DNA binding activity on consensus sequences containing a 5'methylcytosine within the FHL consensus motif. The left panel shows representative gels with the results of quantification plotted on the right.

We tested this model genome-wide using FOXN1 binding data generated in mice(5). After controlling for the presence of enhancer elements, there was a significant enrichment of FOXN1 binding to FOXN1 recognition motifs within CpG islands (4.2-fold, p < 0.0001) but none for FOXN1 motifs located outside of CpG islands (0.9-fold, p = 0.1). This provides support for the above model in which FOXN1 binds to its cognate motifs in the context of a CpG island.

Thus there is the distinct possibility that FOXN1 may be under epigenetic regulation, with changes in the methylation of promoter sequences regulating FOXN1 binding. Global DNA methylation patterns are known to change as a function of age(28), with the general pattern of genome wide hypomethylation and promoter-specific hypermethylation(29). The consequent alteration could lead to a gradual loss of FOXN1 binding to DNA and may thus be a contributing factor to the phenomenon of thymic involution and thus immunosenescence.

## Acknowledgements

The authors would like to thank Diamond Light Source for beamtime (proposal mx15433) and for assistance by staff of beamlines I03 and I04.

The SGC is a registered charity (number 1097737) that receives funds from AbbVie, Bayer Pharma AG, Boehringer Ingelheim, Canada Foundation for Innovation, Eshelman Institute for Innovation, Genome Canada, Innovative Medicines Initiative (EU/EFPIA) [ULTRA-DD grant no. 115766], Janssen, Merck KGaA Darmstadt Germany, MSD, Novartis Pharma AG, Ontario Ministry of Economic Development and Innovation, Pfizer, São Paulo Research Foundation-FAPESP, Takeda, and Wellcome [106169/ZZ14/Z]. AEH is supported by a NIHR Clinical Lectureship.

